# A highly attenuated SARS-CoV-2 related pangolin coronavirus variant has a 104nt deletion at the 3′-terminus untranslated region

**DOI:** 10.1101/2022.02.07.479352

**Authors:** Shanshan Lu, Shengdong Luo, Chang Liu, Muli Li, Xiaoping An, Mengzhe Li, Jun Hou, Huahao Fan, Panyong Mao, Yigang Tong, Lihua Song

**Affiliations:** Beijing Advanced Innovation Center for Soft Matter Science and Engineering, College of Life Science and Technology, Beijing University of Chemical Technology, Beijing, China; Department of Infectious Diseases, Fifth Medical Center of Chinese PLA General Hospital, Beijing, China; Department of Pathology, National Cancer Center/National Clinical Research Center for Cancer/Cancer Hospital & Shenzhen Hospital, Chinese Academy of Medical Sciences and Peking Union Medical College, Shenzhen, China

**Author notes:** These authors contributed equally: Shanshan Lu, Shengdong Luo. Corresponding authors (LS); (YT); (PM); (HF).

## Abstract

SARS-CoV-2 related coronaviruses (SARS-CoV-2r) from Guangdong and Guangxi pangolins have been implicated in the emergence of SARS-CoV-2 and future pandemics. We previously reported the culture of a SARS-CoV-2r GX_P2V from Guangxi pangolins. Here we report the GX_P2V isolate rapidly adapted to Vero cells by acquiring two genomic mutations: an alanine to valine substitution in the nucleoprotein and a 104-nucleotide deletion in the hypervariable region (HVR) of the 3’-terminus untranslated region (3’-UTR). We further report the characterization of the GX_P2V variant in *in vitro* and *in vivo* infection models. In cultured Vero and BGM cells, the GX_P2V variant produced minimal cell damage and small plaques. The GX_P2V variant infected golden hamsters and BALB/c mice but was highly attenuated. Golden hamsters infected intranasally had a short duration of productive infection. These productive infections induced neutralizing antibodies against pseudoviruses of GX_P2V and SARS-CoV-2. Collectively, our data show that the GX_P2V variant is highly attenuated in *in vitro* and *in vivo* infection models. Attenuation of the variant is likely due to the 104-nt deletion in the HVR in the 3’-UTR. This study furthers our understanding of pangolin coronaviruses pathogenesis and provides novel insights for the design of live attenuated vaccines against SARS-CoV-2.

## Introduction

The COVID-19 pandemic caused by SARS-CoV-2 (severe acute respiratory syndrome coronavirus 2) is producing unprecedented damages around the globe^1–3^. As of 20^th^ December 2021, the World Health Organization has reported over 273 million cases with over 5.3 million death cases. Investigation of the origin of the virus has not identified its direct ancestral viruses, but has led to the discovery of many SARS-CoV-2 related coronaviruses (SARS-CoV-2r) in both bats and pangolins^4–9^. Bat-derived coronaviruses found in Laos are the closest relatives of SARS-CoV-2 reported to date^10^. Despite the finding of these SARS-CoV-2r coronaviruses, very few of them have been cultured, thus their biology and pathogenicity remain largely unknown.

While bat viruses are the closest relative of SARS-CoV-2, pangolin coronaviruses likely also play an important role in the origin and evolution of SARS-CoV-2. Two SARS-CoV-2r pangolin coronaviruses GD/2019 and GX/2017 have been identified^6,7^. The GD/2019 strain has a receptor binding domain (RBD) that shares high homology (97.4%) with that of SARS-CoV-2, which leads to the possibility that the origin of the SARS-CoV-2 RBD is a result of recombination between bat and pangolin coronaviruses^7^. The GX/2017 strain has an RBD that shares 86.8% identity with that of SARS-CoV-2^6^. A recent study shows that the spike protein of GX/2017 has comparable binding capabilities to human ACE2 and mediating cell entry when compared to the SARS-CoV-2 spike^11^. Because of the high similarity of RBDs of pangolin CoVs binding to ACE2 receptors between humans and pangolins, pangolin coronaviruses likely provide a repertoire of CoVs for future pandemics.

We previously reported the culture of a pangolin coronavirus GX_P2V^6^, which shares an identical spike protein with other pangolin GX/2017 strains. The genome of coronavirus GX_P2V (GenBank accession number MT072864) was determined by using RNAs from the first passage of the GX_P2V sample, not a serially passaged isolate^6^. This GX_P2V genome has 29,795 nucleotides, including ten unidentified nucleotides and two unknown termini. Considering the similarities in genetics, cell infectivity, and host tropism between this pangolin coronavirus and SARS-CoV-2, we sought to determine the complete genome of the GX_P2V isolate and investigate its biology and pathogenicity in animal models, which may help in understanding the evolution and pathogenicity of SARS-CoV-2 related viruses.

Here, we report that, compared to the original GX_P2V sample, the GX_P2V isolate is a variant with two mutations: one nonsynonymous mutation in the final amino acid codon of the nucleoprotein gene and a 104-nucleotide deletion in the hypervariable region (HVR) of the 3’-terminus untranslated region (3’-UTR). We then describe the characterization of the GX_P2V variant’s growth in two monkey cell lines and two small animal models. We found that the GX_P2V variant has the capability to infect but is highly attenuated in all four of the tested infection models. The attenuation of this pangolin coronavirus isolate is likely due to the 104-nt deletion at 3’-UTR. A short duration of productive infection in golden hamsters induced neutralizing antibodies against pseudoviruses of GX_P2V and SARS-CoV-2. This work advances our understanding of pangolin coronaviruses pathogenesis and provides important insights for the design of live attenuated vaccines against SARS-CoV-2.

## Results

### Genomic determination of pangolin coronavirus GX_P2V isolate

Total RNAs from the eighth viral passage were used for Illumina pair-end sequencing. In total, 8,482,736 reads were mapped to the previous genome of GX_P2V using Bowtie 2. The resulting viral reads were then assembled *de novo* by Trinity. The viral genomic contig has 29,718 nucleotides including eight consecutive adenine residues at the 3′-terminus. The 5′-terminus of the viral genome was determined by 5′/3′ RACE kits (TaKaRa) and the whole genome sequence (29,729 nt) was deposited in GenBank (accession number MW532698).

### Genomic comparison between the GX_P2V sample and the coronavirus GX_P2V isolate revealed two mutations in the passaged viral isolate

We next compared the sequence differences between the first passage of the GX_P2V sample and the eighth passage of the GX_P2V isolate. Compared to the single cell-passaged sample, the multiply passaged GX_P2V isolate has complete 5′/3′-terminus sequences and two mutations: a nonsynonymous mutation in the final amino acid codon of the nucleoprotein gene (GCU→GUU) and a 104-nucleotide deletion in the hypervariable region (HVR) of the 3’-terminus untranslated region (3’-UTR) (Fig. 1). Thus, the multiply passaged GX_P2V isolate is a cell culture adapted variant. The result of the single-nucleotide mutation is an alanine to valine change at the carboxyl terminal of the nucleoprotein. We then asked whether these two mutations occurred in the first passage of the GX_P2V in Vero cells. Among the sequencing reads of the GX_P2V sample (SRR11093271), only one read contained the nucleoprotein single-nucleotide mutation while no reads contained the HVR deletion. In comparison, among the sequencing reads of the multiply passaged isolate, all reads covering these two mutated regions contain the identified mutations. These data suggest that these two identified regions have strong selection pressures in cell culture and selection occurred rapidly following *in vitro* growth.

**Fig. 1.**
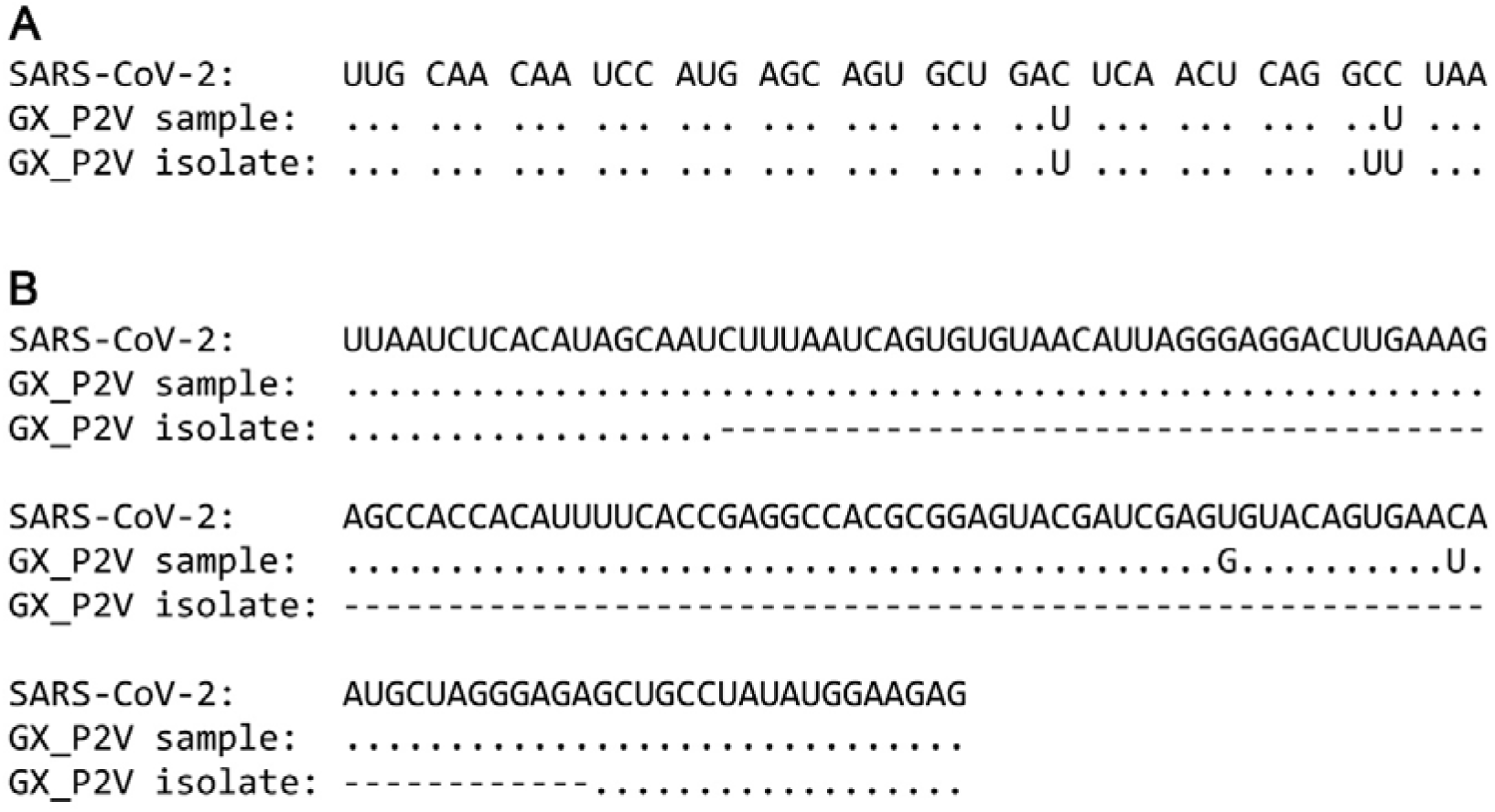
Sequence comparisons between SARS-CoV-2, the original GX_P2V sample and the pangolin coronavirus GX_P2V isolate. The GX_P2V isolate is a variant with two interesting mutations: one nonsynonymous mutation (C to U) in the final amino acid codon of the nucleoprotein and one 104-nt deletion in the 3′-UTR. **a** Alignment of the 3′-terminus sequences of the nucleoprotein genes. **b** Alignment of the partial 3′-UTR sequences. Dots represent residues that are identical to those in SARS-CoV-2 and hyphens represent nucleotide deletions.

The temporal occurrence and selective factor(s) for these mutations are unknown. It is plausible that both mutations increased the efficiency of viral replication. The spontaneous 104-nt deletion within the viral 3’-UTR is of special interest. According to the current 3’-UTR model^12^, the HVR region of the GX_P2V sample is 166-nt long -ranging from 29,611 to 29,776 based on the deposited sequence data (GenBank accession number MT072864). The deletion region ranges from 29,617 to 29,720, covering 62.7% of the entire HVR region. The HVR region is absolutely maintained in all coronavirus clinical samples. Based on our knowledge, this is the first report of a spontaneous deletion in the HVR region.

### Growth of the pangolin coronavirus GX_P2V variant yields mild cell damage and small plaques in non-human primate cells

The GX_P2V isolate was cultured from a mixture of pangolin lung and small intestine samples by using Vero E6 cells^6^. It has been shown that the virus only partially destroyes Vero E6 cell monolayer five days post-infection^6^. In cells infected with the virus for five days, the cytopathic effects manifested as cell degeneration and cell lysis, with significant portion of cell rounding but limited cell detachment. We next systematically investigated the growth of the GX_P2V variant in two non-human primate cells: Vero and BGM. The BGM cell line was selected based on the fact that BGM is susceptible to SARS-CoV and the pangolin viruses GX/2017 utilize ACE2 as the cell receptor for viral attachment and entry^11,13^.

BGM and Vero cells were infected with the GX_P2V variant at a multiplicity of infection (MOI) of 0.01. Growth of GX_P2V induced similar cytopathic effects in both Vero and BGM cells (Fig. 2a). No or limited cell degeneration can be observed at 24 hours postinfection. More evident cell degeneration and cell rounding appeared at 48 hours postinfection. Cell lysis and detachment started around 72 hours postinfection. These cytopathic effects were directly related to viral replications, which was confirmed by indirect immunofluorescence (Fig. 2b). Consistent with the observed slow rate of cell damages, the GX_P2V variant produced small plaques at five days postinfection (Fig. 2c, d). In comparison, SARS-CoV-2 produced substantial cell damages and large plaques in Vero cells at two days postinfection^4,14^. Compared to reported SARS-CoV-2 data, the GX_P2V variant is highly attenuated in causing cell damage and plaque formation.

**Fig. 2.**
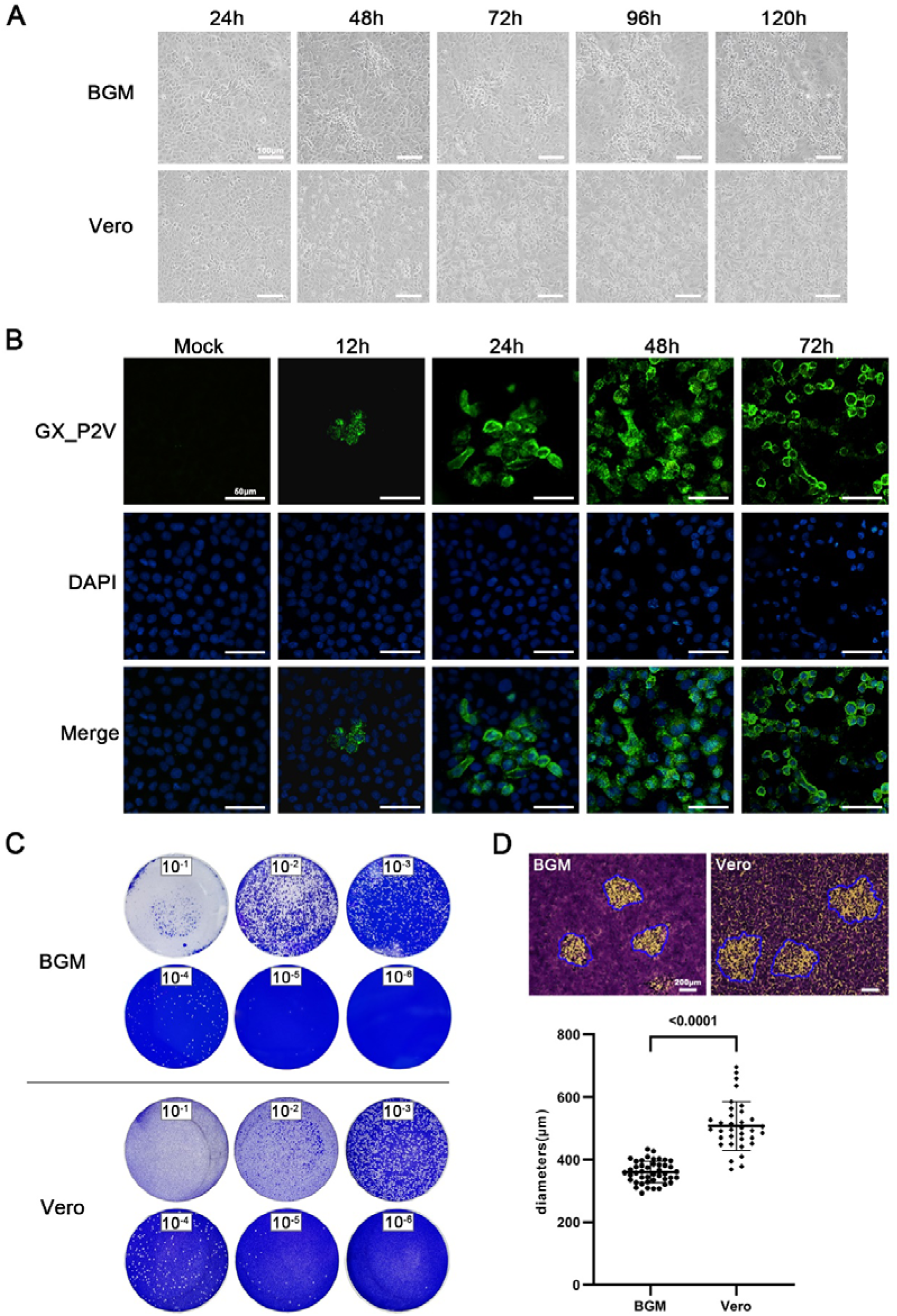
Growth of the pangolin coronavirus GX_P2V variant in two non-human primate cells: BGM and Vero. **a** BGM and Vero cells infected with the GX_P2V variant show similar cytopathic effects: limited cell degeneration at 24 h postinfection, and more evident cell degeneration and cell rounding after 48 h postinfection. Scale bars, 100 μm. **b** The replication of the GX_P2V variant in BGM cells at different time points was detected by immunofluorescence assays with anti-sera from GX_P2V-infected hamsters. Nuclei were stained with DAPI (blue). Scale bars, 50 μm. **c** The GX_P2V variant forms small plaques in BGM and Vero cells five days postinfection. The small white dots represent plaques. **d** The diameter of plaques (blue circle) was measured by using Image Pro plus 6.0 software. The GX_P2V variant forms bigger plaques in Vero than in BGM (*p* < 0.0001).

The growth kinetics of the GX_P2V variant in both BGM and Vero cells were measured by quantifying the viral RNA copies and viral titers in the culture supernatants by using qPCR and viral titration, respectively (Fig. 3). Cells were infected with a MOI of 0.01. Supernatants were collected at indicated time points and were used for quantification assays. In both cells, the viral RNA copies reached the peak at 72 h.p.i. (Fig. 3a) and the peaks of viral titers were at 48 h.p.i. (Fig. 3b), though only limited cell degenerations could be observed at this time point. The viral titers at 48 h.p.i. were as high as 10^7^ TCID_50_/mL -a titer number that is comparable to the titer of SARS-CoV-2 at this time point^14^. The growth curves of the GX_P2V variant in both cells are similar, but do have significant differences at 24 and 120 h.p.i. (Fig. 3b). Higher viral titers were detected in Vero cells at 120 h.p.i. (*p* < 0.001). Consistent with this difference, Vero cells were more supportive of viral plaque assays, as they have more cell lysis starting at 72 h.p.i. and produced larger viral plaques at 120 h.p.i. (0.36 ± 0.04 mm v.s. 0.51 ± 0.08 mm, *p* < 0.0001) (Fig. 2a, d).

**Fig. 3.**
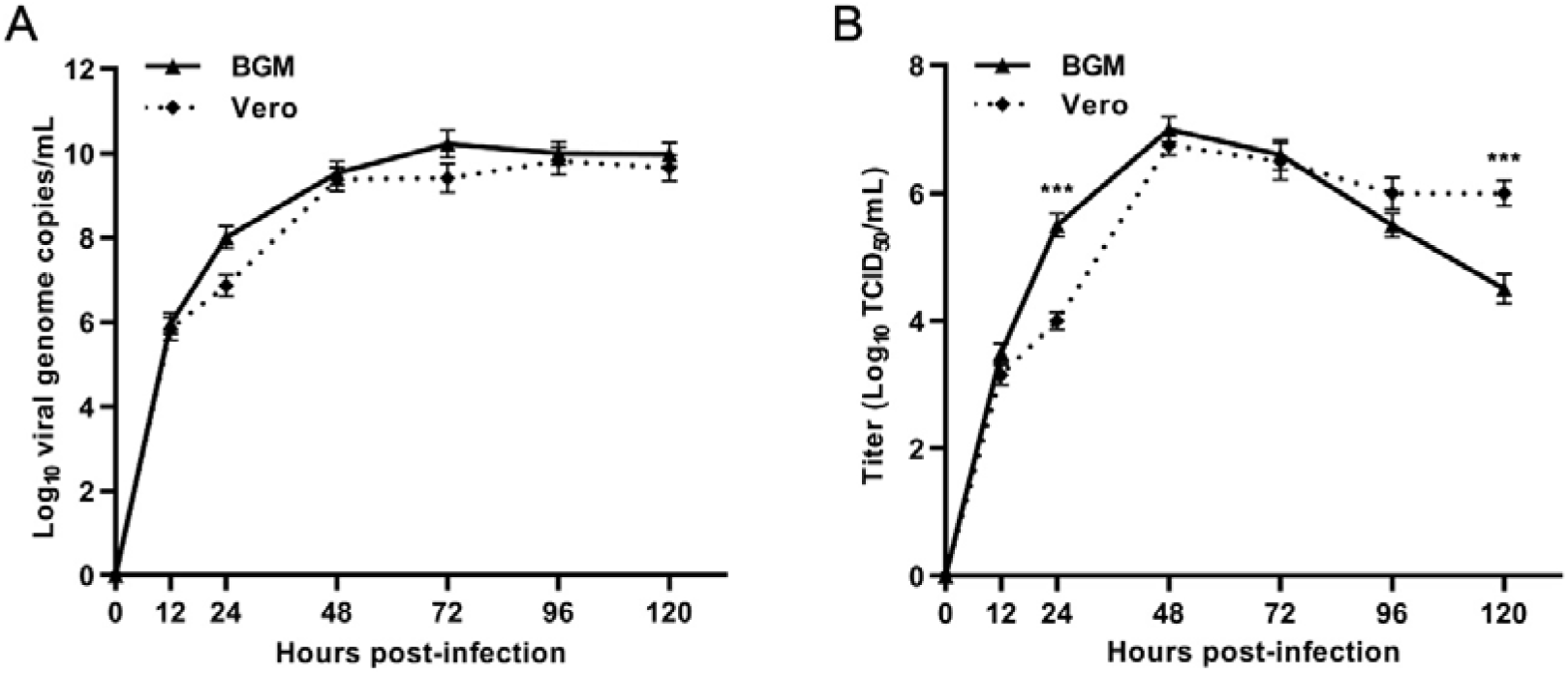
Replication kinetics of the GX_P2V variant in BGM cells (solid line) and Vero cells (dotted line). Results are expressed as the mean of three biological replicates. **a** Viral RNA copies were determined by qRT-PCR. **b** Viral titers were titrated by using a standard TCID_50_ assay. The two cells appear to produce different viral titers at 24 and 120 h postinfection (^***^*p* < 0.001).

### The pangolin coronavirus GX_P2V variant is highly attenuated in the golden hamster intranasal infection model

Studies on SARS-CoV-2 pathogenesis provide animal models for investigating the pathogenicity of the GX_P2V variant^15–17^, as its spike protein has similar capabilities of binding to human ACE2^11^. The golden hamster model is especially susceptible to SARS-CoV-2. SARS-CoV-2 administered by intranasal route can replicate efficiently in the lungs of hamsters, causing severe pathological lung lesions^16^. These severe lung injuries shared characteristics with SARS-CoV-2-infected human lungs, including severe, bilateral, peripherally distributed, multilobular ground glass opacity, and regions of lung consolidation^16,17^. We next investigated the replicative ability and pathogenesis of the GX_P2V variant in golden hamsters.

Three groups of hamsters were intranasally infected with mock, 1×10^4^ and 1×10^5^ TCID_50_ of the GX_P2V variant, respectively. Clinical signs were recorded and samples were collected periodically throughout the 14-day experiment (Fig. 4a). The infection outcomes are clearly contrast to those described in the SARS-CoV-2 infected hamsters. Hamsters infected with the GX_P2V variant had similar body weight increases to the mock infected hamsters and no apparent clinical symptoms in the two week observative period (Fig. 4b). Consitent with the absence of clinical manifestations, there was only a slight hyperemia noted on the surface of lungs following infection with the GX_P2V variant at 2 and 5 d.p.i. (Fig. 4c). Moreover, no signs of bronchopneumonia were found in the tracheas of infected hamsters (Fig. 4c).

**Fig. 4.**
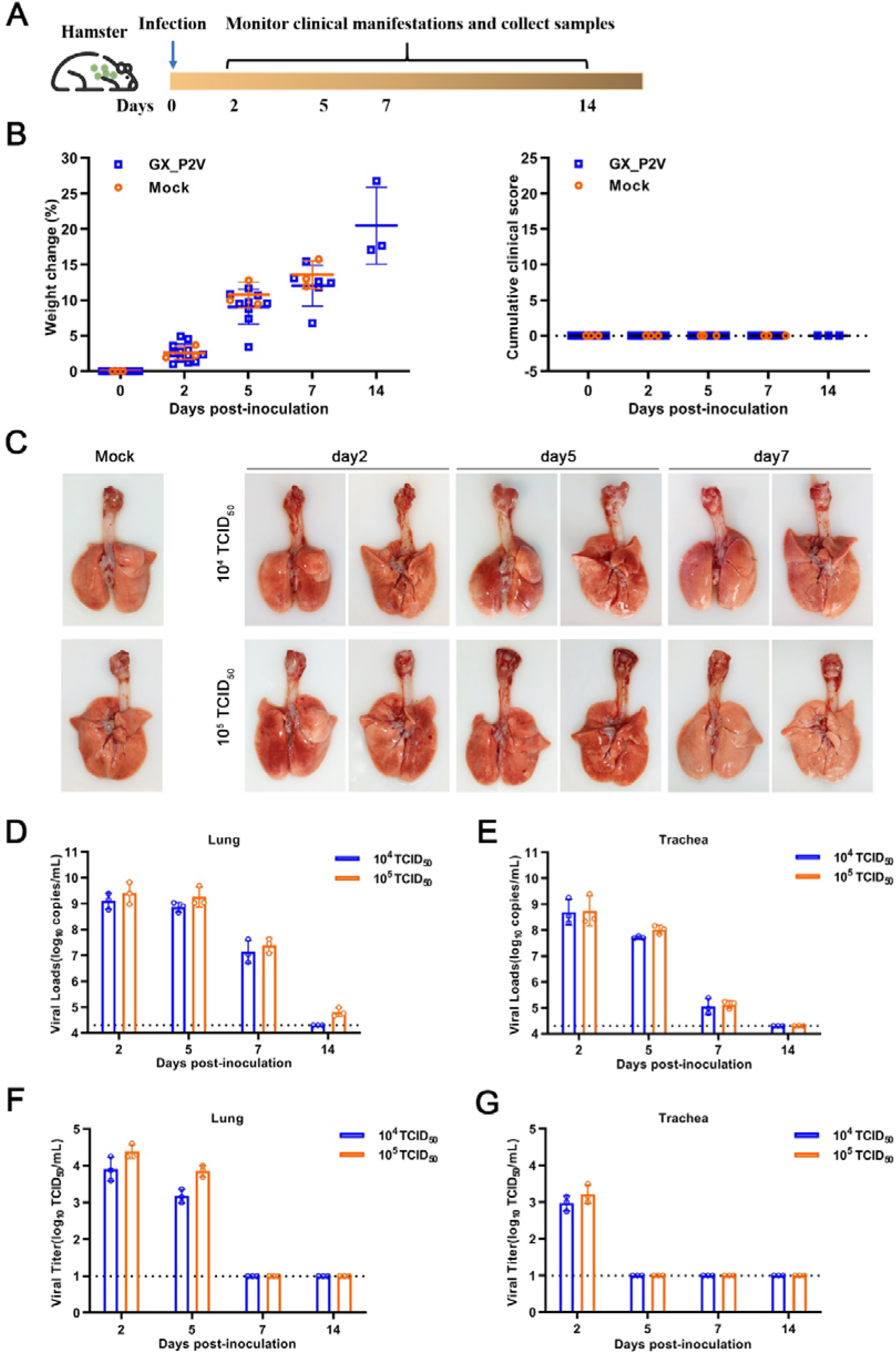
Intranasal infection of golden hamsters shows the GX_P2V variant had a short duration of viral replication and caused no gross pathology in the respiratory tracts. **a** Scheme of intranasal infection. **b** Both mock and GX_P2V-infected hamsters had normal body weight increases and no apparent clinical symptoms in the two-week observational period. Individual data points represent percent weight changes compared to day zero, or cumulative clinical score. **c** Representative images of lungs and tracheas from mock hamsters, 10^4^, and 10^5^ TCID_50_ infected hamsters at indicated time points. All organs had no gross pathology. **d-g** Viral RNA loads (**d, e**) and viral titers (**f, g**) in homogenate supernatants of lungs (**d, f**) and tracheas (**e, g**) of GX_P2V-infected hamsters at different days post-inoculation were determined by qRT-PCR and the TCID_50_ assay, respectively. The detection limit is shown by the dotted line. Error bars represent means±SD.

To monitor the viral replication and pathogenesis, lungs and tracheas of hamsters were collected at indicated time points. Viral RNA copies and viral titers were measured. In both organs, viral RNA copies peaked at two d.p.i. and gradually decreased for the remaining says (Fig. 4d, e). The RNA copy numbers at two d.p.i. were larger than the original inoculum numbers, demonstratng replication in both lungs and tracheas. However, viral titers were only detected in early stages post-infection: at two and five d.p.i. in lungs and day two in tracheas (Fig. 4f, g). This suggests a shorter duration of viral replication in both organs, but especially in tracheas.

Hematoxylin and eosin staining was performed to determine the histopathological changes in both lungs and tracheas. Histopathologically, the tissues from hamsters of the 1×10^4^ TCID_50_ group were comparable to sham infected controls (Fig. 5a). Infection with 1×10^5^ TCID_50_ of the GX_P2V variant resulted in a local infiltration with very few inflammatory cells at five d.p.i., and none at other time points (Fig. 5b). Compared to the mock group, no pathological lesions were found in tracheas of all GX_P2V variant infected hamsters over the entire observation period (Fig. 5c). Thus, the GX_P2V isolate did not induce potentially damaging inflammatory changes in lungs or tracheas of either infected groups. The lack of detectable pathology and short duration of viral replication in infected hamsters suggests that the GX_P2V variant has a replicative deficiency in hamsters.

**Fig. 5.**
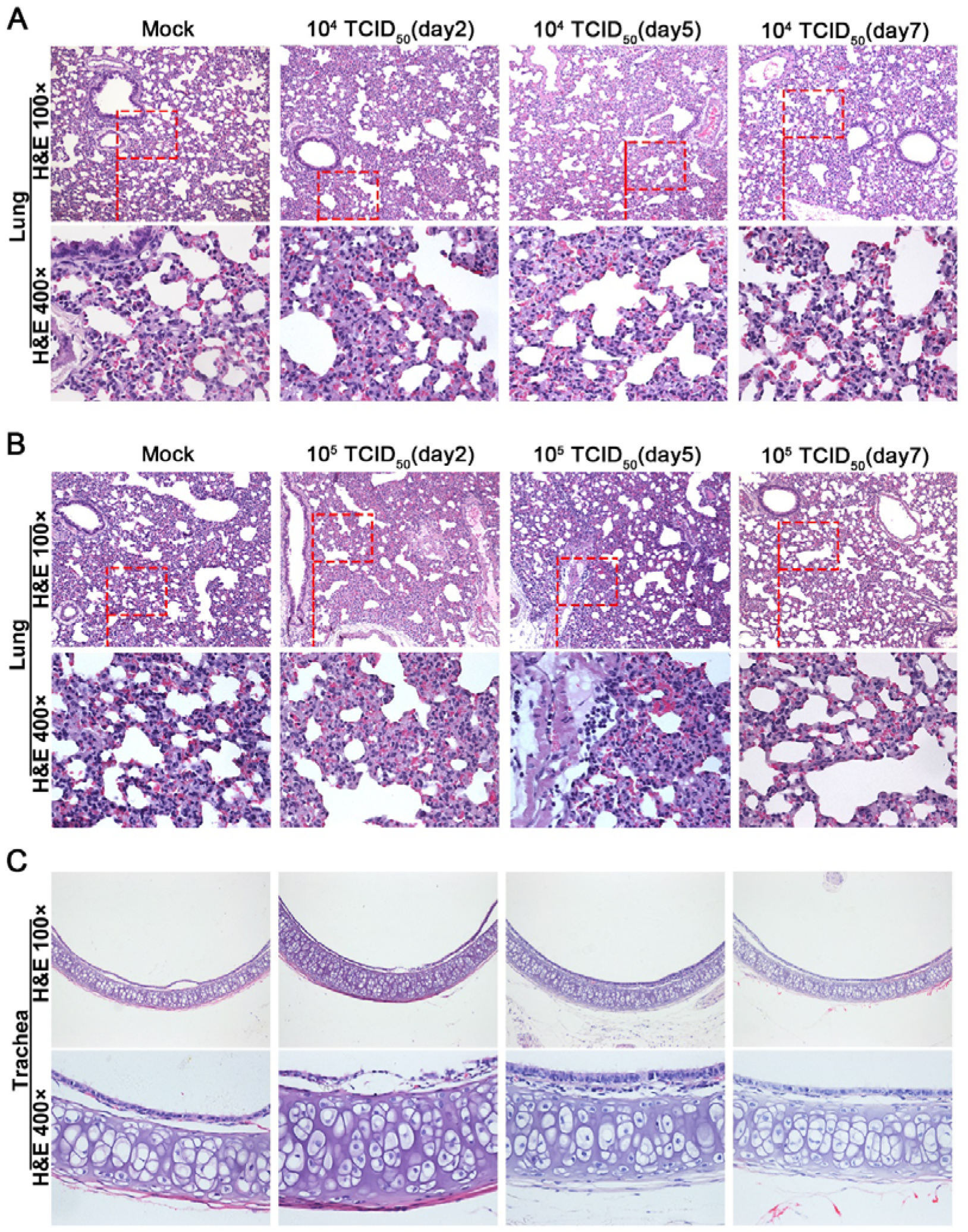
Histopathological analyses of lung and trachea tissues from intranasally infected hamsters. H&E 400 × are the magnifications of the region in the corresponding red box in H&E 100 ×. **a** 10^4^ TCID_50_ infected group (n = 3), no pathological changes were observed in the lungs. **b** 10^5^ TCID_50_ infected group (n = 3), lungs had no tissue lesions. Few inflammatory cells of lymphocytes and neutrophils associated with blood vessels were observed at five days postinfection but not at other time points. **c** Representative images show that tracheas of both infection dose groups had no significant histopathological changes at all time points post-infection.

### Intragastrical inoculation established limited infections in respiratory tracts but no detectable infection in the gastrointestinal tract

Human gastrointestinal (GI) infection by SARS-CoV-2 has been implied, as SARS-CoV-2 RNA can be frequently detected in the feces of COVID-19 patients^18^. In the golden hamster model, oral inoculation can establish a SARS-CoV-2 infection with an infection pattern that is consistent with mild COVID-19^19^. In the above described intranasal infection of golden hamsters with the GX_P2V variant, viral RNA sheddings were detected in tongues and feces only at the earliest times post inoculation (Figue. 6a), which is different than the continuous RNA sheddings in COVID-19 patients. We performed intragastrical inoculation of the GX_P2V variant in golden hamsters to further assess viral pathogenicity. Golden hamsters were intragastrically infected with 10^5^ TCID_50_ of the GX_P2V variant and samples were collected periodically (Fig. 6b). All hamsters had consecutive weight increases (Fig. 6c) and no apparent clinical signs were observed (Fig. 6d). Limited viral RNAs were detected in lungs but not in other organs or samples (stomachs, duodenums, colons, tongues, trachea, esophaguses, small intestines and feces) (Fig. 6e). Thus, the virus shedding of GX_P2V in golden hamsters differs from the chronic virus shedding from the oral cavity and feces following oral inoculation of SARS-CoV-2^19^. Predictably, no histopathological changes were detected in lungs of intragastrically infected hamsters (Fig. 6f). Similar to previous reports with SARS-CoV-2 in golden hamsters^19^, we failed to detect infectious virus in fecal samples from either intranasal or intragastrical infections (Fig. 6g). Thus, in golden hamsters, the GX_P2V variant has limited capability of establishing infections of the gastrointestinal tract or when introduced by the intranasal route.

**Fig. 6.**
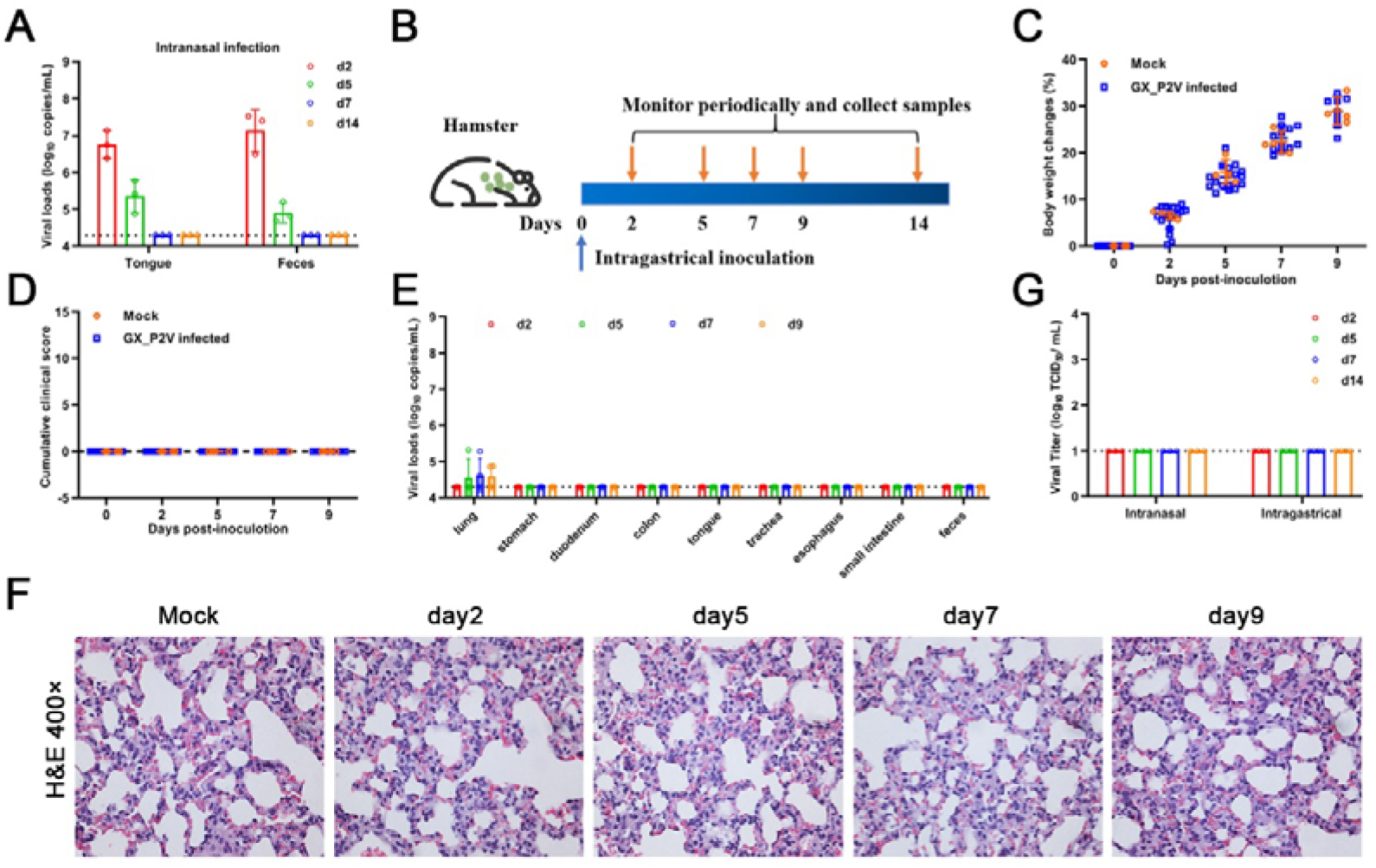
Intragastrical inoculation of the GX_P2V variant in golden hamsters established limited infection in the respiratory tract. **a** Viral RNAs were detected in samples of tongues and feces from the above golden hamsters with intranasal infection, suggesting established infections in the gastrointestinal tract. **b** Scheme of intragastrical infection. Samples of tongues and feces were collected at indicated time points. **c** Body weights of all golden hamsters had continuous increases. **d** No clinical symptoms of infection were identified. **e** Limited viral RNAs in the lungs, but not in the stomach, colon or faeces were detected by qPCR. **f** No histopathological changes were identified in lung tissues of mock and GX_P2V-inoculated golden hamsters. **g** Viruses were not detected in feces from golden hamsters following intranasal or intragastrical inoculation.

### The GX_P2V variant does replicate but produce no infectious virus in young and aged BALB/c mice

SARS-CoV-2 can rapidly adapt in aged BALB/c mice and induce typical pneumonia^20,21^. Therefore we next investigated the pathogenicity of the GX_P2V variant in both young and aged BALB/c mice. BALB/c mice were intranasally infected with 10^5^ TCID_50_ of the GX_P2V variant and euthanized to collect samples at specific days postinfection (Fig. 7a). Following inoculation, no apparent clinical signs were observed in any inoculated mice. Noteably, the body weights of young BALB/c mice continued to increase (Fig. 7b) and no gross pathological changes on the lung surfaces of inoculated mice were detected (Fig. 7c). The viral RNAs in the lung of mice were rapidly cleared (Fig. 7d), and aged mice had an even faster rate of viral clearance. This is in contrast to the slower rate of viral clearance in aged mice inoculated with SARS-CoV-2^20^. Histopathological analyses suggest that lymphocytes play an important role in the rapid viral clearing (Fig. 7e, f). Lungs of the inoculated young mice had local thickened alveolar septa and lymphocyte aggregations at one d.p.i., then rapidly returned to normal at three d.p.i. (Fig. 7e). Comparatively, the lungs of the inoculated aged mice showed multifocal abundant lymphocytes and some macrophages present up to day 3 d.p.i. (Fig. 7f). Furthermore, we failed to detect any virus in lungs, despite detection of high viral RNA copies at the early infection time points (Fig. 7f). Altogether, our data indicate that the GX_P2V variant can infect but has a replication deficiency in BALB/c mice.

**Fig. 7.**
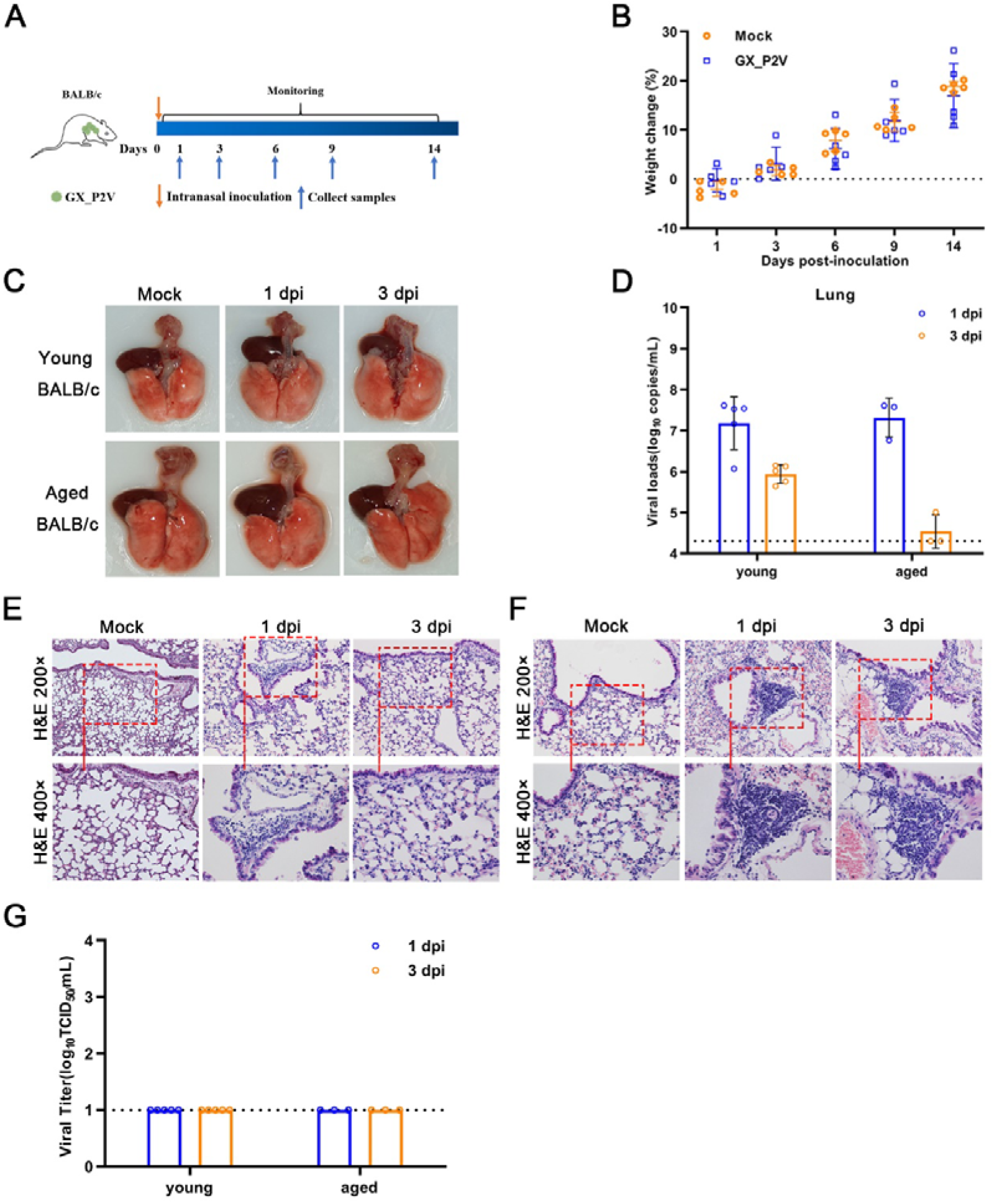
Intranasal inoculation of the GX_P2V variant in both young and aged BALB/c mice. **a** Scheme of intranasal infection. Young and aged BALB/c mice were intranasally infected with mock or 10^5^ TCID_50_ of the GX_P2V variant. Various infection outcomes and tissue samples were collected at indicated time points. **b** Young mice had constant body weight increases after infection. **c** Intranasal inoculation of GX_P2V in young and aged BALB/c mice produce no detectable gross pathology. **d** Viral RNA loads in infected lungs of both young and aged BALB/c mice were determined by qPCR. **e-f** Representative H&E images of infected lungs from young (**e**) and aged (**f**) BALB/c mice. **g** No viral titers were detected in infected lungs from both young and aged BALB/c mice.

### Productive infections in golden hamsters induced neutralizing antibodies against both GX_P2V and SARS-CoV-2

The coronavirus GX_P2V has high genomic homology with SARS-CoV-2^6^ The spike protein and the receptor binding domain of GX_P2V share 92.4% and 86.8% amino acid identity with those of SARS-CoV-2, respectively. The attenuating nature of the GX_P2V variant, combined with its similarity to SARS-CoV-2, led us to propose that the GX_P2V variant is a potential live vaccine candidate. To address, we analyzed the humoral responses against GX_P2V in infected animals and evaluated cross-reactive antigenicity of the two spike proteins from GX_P2V and SARS-CoV-2.

Cells transfected with the full-length genes of the two spike proteins were used to perform immunofluorescence assays with sera from both convalescent COVID-19 patients and intranasally infected golden hamsters (Fig. 8a). The results clearly show that these two spike proteins share strong cross-reactive antigenicity. Commercial SARS-CoV-2 S1 antibody assay kits can be used to titrate the humoral responses against GX_P2V in hamsters (Fig. 8b). The intranasally infected hamsters have a strong humoral response to the spike protein. Such strong immune responses correlate with the productive infections in intranasally infected hamsters. In contrast, low titer or no antibodies against the spike protein were detected in intragastrically infected hamsters or intranasally infected BALB/c mice (Fig. 8c).

**Fig. 8.**
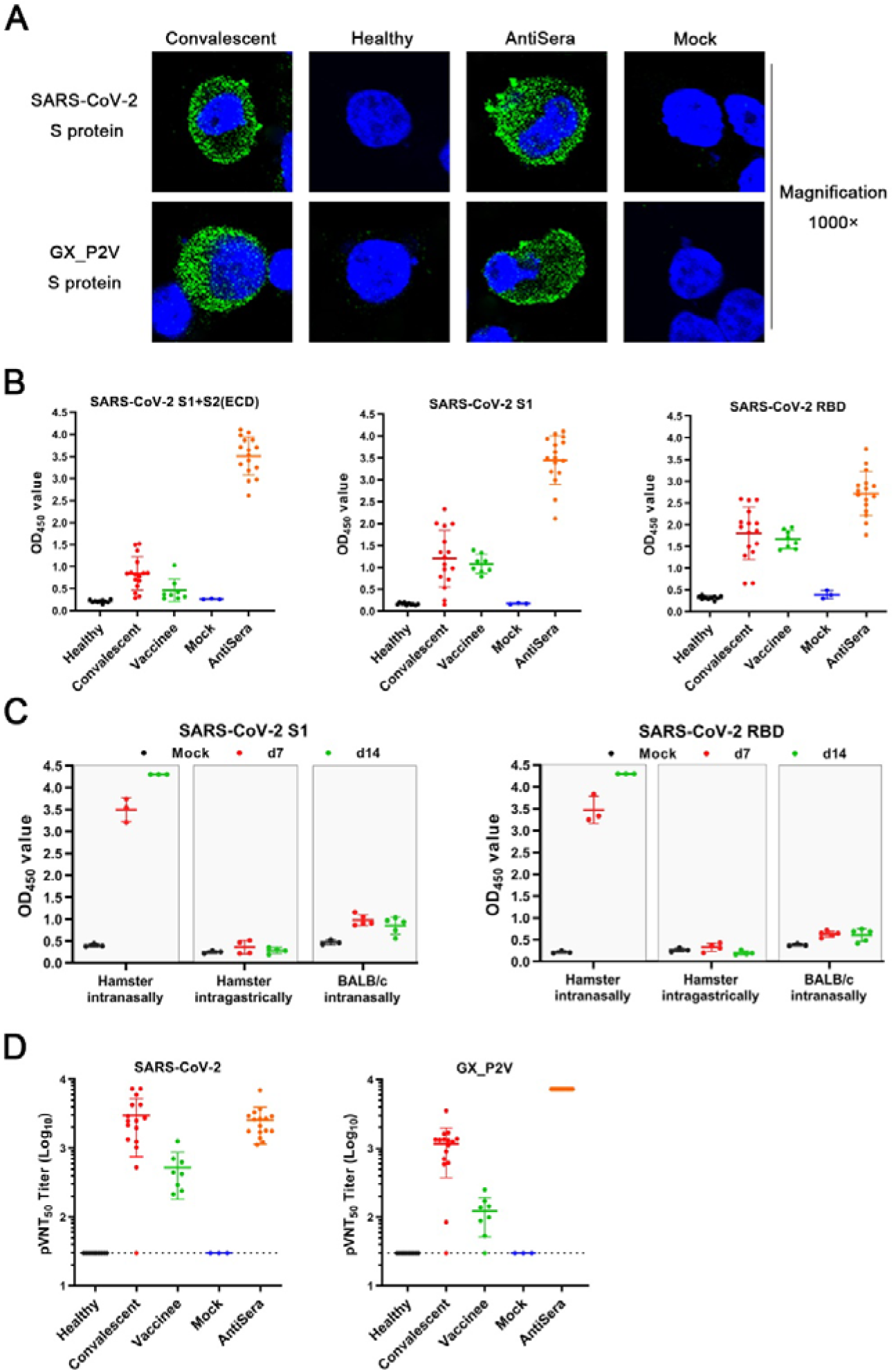
Serological cross-reactivity and cross-neutralization between GX_P2V and SARS-CoV-2. **a** Immunofluorescence assays with 293T cell expressing spike proteins (green) of SARS-CoV-2 and GX_P2V, using sera from convalescent COVID patients, healthy donors, and GX_P2V-infected golden hamsters. Nuclei were stained with DAPI (blue). **b** Three commercial ELISA kits for detecting antibodies against SARS-CoV-2 S1+S2 (ECD), S1, and RBD were used for measuring antibodies in sera from healthy donors (n = 10), convalescent COVID patients (n = 16), vaccinees of inactivated COVID-19 vaccines (n = 8), mock-infected golden hamsters (n = 3), and GX_P2V-infected golden hamsters (n = 16), respectively. **c** Cross-reacting antibodies against SARS-CoV-2 S1 and RBD in sera from GX_P2V-infected golden hamsters and BALB/c mice were measured by using the aforementioned ELISA kits. No cross-reacting antibodies against SARS-CoV-2 S1 and RBD were detected in sera from intragastrically infected golden hamsters or intranasally infected BALB/c mice. **d** Neutralizing antibodies against two pseudoviruses, SARS-CoV-2 and GX_P2V, were detected in sera as described in figure 8b. The error bars indicate means±SD.

SARS-CoV-2 neutralizing antibodies play an important role for the protection against COVID-19. We analyzed the neutralizing antibodies in sera from COVID-19 convalescent patients, vaccinees of inactivated COVID-19, and intranasally infected hamsters by using pseudoviruses of GX_P2V and SARS-CoV-2 (Fig. 8d). All tested positive sera had variable titers of neutralizing antibodies against both pseudoviruses. When compared to sera from convalescent patients, the hamster sera had a similar titer of neutralizing antibodies against SARS-CoV-2, although the RBDs of GX_P2V and SARS-CoC-2 share only 86.8% amino acid identity and share a single critical receptor-binding residue^6^. This finding supports the rational for the design of pan-SARS-CoV-2r vaccines.

## Discussion

The devastating force of COVID-19 intensified the world’s fear over SARS-CoV-2 and its related coronaviruses. The search and characterization of SARS-CoV-2 related coronaviruses is an area of intense interest. In this study, we investigated the genome, biology, pathogenicity, and cross-reactive antibody response of a SARS-CoV-2 related pangolin coronavirus GX_P2V variant. We found that this GX_P2V variant is a 3’-UTR shortened mutant derived from the original GX_P2V sample. The GX_P2V variant was highly attenuated in two cell cultures, golden hamsters, and BALB/c mice. Importantly, the antibody elicited by GX_P2V isolate infection in hamsters inhibited SARS-CoV-2 spike-pseudovirus infection. Attenuation and cross-neutralizing activity of the GX_P2V variant provides new insights for our understanding of coronavirus pathogenesis and for developing a live attenuated vaccine for SARS-CoV-2. Unfortunately, the original wild type GX_P2V virus was not isolated thus it could not be comparably studied. Nevetheless, this study represents the first report of characterizing SARS-CoV-2 related coronavirus in animal models.

In cell cultures, attenuation of the GX_P2V variant caused mild cell damage and small plaques. The attenuation is likely independent of viral infectivity, as the spike proteins of GX_P2V and SARS-CoV-2 have comparable capabilities of mediating cell attachment and entry^11^. It is also likely viral replication and egress were unaffected, as titers of live GX_P2V in the supernatants reach similar levels at 48 h.p.i. as SARS-CoV-2^14,22^. How could GX_P2V produce these attenuating phenotypes while having growth capabilities similar to those of SARS-CoV-2? Certain mutations in the spike protein have been linked with viral plaque sizes in SARS-CoV-2^22^. Therefore, it is possible the spike protein of GX_P2V is less effective in host cell interactions than that of SARS-CoV-2.

The attenuation of the GX_P2V variant in animal models can be explained by two aspects of our findings. The first is the 104-nucleotide deletion in the HVR of the 3’-UTR of the GX_P2V variant. In the mouse hepatitis virus (MHV) model, the association between HVR mutations and viral pathogenicity has been comprehensively investigated^23^. MHV with a deletion of the entire HVR had minimal change on viral growth in cell cultures, but was highly attenuated in mice, causing no signs of clinical disease^23^. The HVR of GX_P2V likely also plays a significant role in viral pathogenesis. Another aspect is the lower inner shell disorder in GX_P2V as predicted by measuring the percentage of protein intrinsic disorder of the nucleoprotein^24^. Goh et al^24^ predicted that the wild type GX_P2V virus was an attenuated SARS-CoV-2 vaccine strain even though the virus was initially identified in dead pangolins.

The golden hamsters can be infected by both SARS-CoV and SARS-CoV-2, but disease outcomes varied by different strains and different infection routes^15,19,25^. Hamsters intranasally inoculated with a SARS-CoV Urbani strain had robust viral replication, with significant pathology in the respiratory tract, but without weight loss or evidence of disease^25^. A follow-up study reported a lethal SARS-CoV Frk-1 strain, which differs from the Urbani strain by an L1148F substitution in the S2 domain^26^. Hamsters intranasally infected with SARS-CoV-2 were found to resemble human infections by multiple groups^15,16,27^. Overall, hamsters intranasally infected with either SARS-CoV or SARS-CoV-2 consistently had similar viral durations and siginificant pathologies in the lungs. While having similar viral loads and similar duration of viral titers in lungs, the GX_P2V variant in hamsters caused no pathology or clincal signs of disease. We surmise the absence of pathology is likely in part due to the low virulence nature of the GX_P2V spike protein.

Live attenuated vaccines are attractive because they can stimulate a strong and durable immune response with both humoral and cellular immunity. The challenge of constructing live attenuated vaccines is the difficulty of adopting acceptable attenuation strategies. Currently, two live attenuated SARS-CoV-2 vaccine candidates generated by using genome recoding have been reported^28,29^. Our data suggest that shortening the 3’-UTR is a novel strategy for making a live attenuated SARS-CoV-2 variant that warrants further investigations.

## EXPERIMENTAL MODEL AND SUBJECT DETAILS

Vero and BGM cells, obtained from the American Type Culture Collection (ATCC) (Manassas, USA), were grown in Roswell Park Memorial Institute (RMPI) 1640 medium (Gibco, USA) supplemented with 10% fetal bovine serum (FBS) (Gibco, USA) and 100 units/mL penicillin-streptomycin (Gibco, USA) at 37°C in 5% CO_2_. Stable hACE2-293T cells were kindly provided by Pro. Zheng Zhang (Shenzhen Third People’s Hospital) and were cultured in DMEM (Gibco, USA) with 10% FBS and 100 units/mL penicillin-streptomycin.

The pangolin coronavirus GX_P2V, originally cultured from the lung-intestine mixed samples of a pangolin captured in anti-smuggling operations in 2017^6^, was passaged in Vero cells. Cell supernatants were harvested and centrifugated (10,000 × *g*) for five minutes, then supernatants were aliquoted and stored at −80°C. Virus titers were determined by using a standard 50% tissue culture infection dose (TCID_50_) assay.

Human serum samples were collected from healthy donors, COVID-19 patients, and vaccinees of two doses of inactivated SARS-CoV-2 vaccine BBIBP-CorV at the Fifth Medical Center of General Hospital of Chinese PLA. All participants signed informed consent forms. The blood serum samples from COVID-19 patients and vaccinees were collected at 21 to 28 days after symptom onset, and two weeks after receiving the second vaccination, respectively. All serum samples were stored at −80°C. This study was approved by the institutional review board of the Fifth Medical Center, General Hospital of Chinese PLA (approval number: 2020027D).

Six-week-old male hamsters, six-week-old male BALB/c and 48-week-old female BALB/c were used in this study. The procedure of animal experiments was approved by the Institutional Animal Care and Use Committee of Fifth Medical Center, General Hospital of Chinese PLA (IACUC-2018-0020).

## METHOD DETAILS

### Next-generation sequencing

The eighth passage of the pangolin coronavirus GX_P2V was partially purified from cell culture supernatants by ultra-centrifugation. Total RNAs were extracted by using the AxyPrep™ body fluid viral DNA/RNA Miniprep kit (Axygen, Cat No. AP-MN-BF-VNA-250, Hangzhou, Zhejiang, China) according to the manufacturer’s instructions. Whole-genome sequencing was performed using next-generation sequencing (NGS) methodology. The cDNA libraries were constructed using a TruSeq RNA library prep kit (Illumina) according to the manufacturer’s instructions and sequenced on an Illumina MiSeq system. Sequence data was deposited in the China National Microbiological Data Center (project accession number NMDC10017729 and sequence accession number: NMDC40002427).

### Assembly and characterization of the coronavirus genome

Raw reads were adaptor- and quality-trimmed with the Fastp program^30^. The clean reads were then mapped to the near complete genome of pangolin coronavirus GX_P2V (GenBank accession number MT072864) using Bowtie 2^31^. The reads that were mapped to the GX_P2V genome were then assembled *de novo* using Trinity with default settings^32^. The assembled contig had a complete 3′-terminus. Then, the 5′-terminus of the viral genome was determined by 5′/3′ RACE kits (TaKaRa). The resulting whole genome sequence of the GX_P2V variant was deposited in GenBank (accession number MW532698). To characterize and map the mutations of the GX_P2V isolate, the viral genomic sequences of the GX_P2V variant and the GX_P2V sample were aligned using ClustalW^33^.

### Plaque assay and cytopathic effect (CPE)

The plaque assay using methylcellulose, a low-viscosity, semi-solid overlay, was performed to determine the plaque phenotype and infectious titers as previously described^34^. Confluent monolayers of BGM and Vero cells seeded in 6-well plates were incubated with ten-fold serially diluted viral stocks ranging from 10^−1^ to 10^−6^ for 2 h at 37°C with gentle rocking every 15 minutes. After viral adsorption, cells were washed twice with phosphate buffer saline (PBS) and overlaid with Minimum Essential Medium (MEM) (Gibco, USA) containing 2% FBS, 1% methylcellulose, and HEPES (20 mM) and incubated at 37°C in 5% CO_2_ incubator. Cytopathic effect (CPE) was observed and photographed daily for 5 days under a light microscopy. At 5 days post-infection (d.p.i.), cells were fixed with 4% paraformaldehyde for 2 h at room temperature, and then stained with 0.2% crystal violet for 20 min. They were gently rinsed under tap water and dried prior to analysis of the plaque phenotype and calculation of virus titers.

### Viral growth kinetics

Confluent monolayers of BGM and Vero cells in 6-well plates were incubated with viral stocks at a multiplicity of infection (MOI) of 0.01 for 2 h at 37°C followed by washing twice with PBS and maintained in RMPI 1640 medium with 2% FBS and HEPES (20 mM) at 37°C in a 5% CO_2_ incubator. Culture supernatants were collected at the indicated time points and stored at −80°C for subsequent viral growth kinetics assay by quantifying the viral RNA copies and viral titers by using qPCR and 50% tissue culture infectious dose (TCID_50_), respectively.

### TCID_50_ assay

BGM cells (2.5 × 10^4^ cells/well) in 96-well plates were incubated together with ten-fold serial dilutions of the cell culture supernatants at 37°C in a 5% CO_2_ incubator. Cytopathic effect was observed at 5 d.p.i. Viral titers were calculated and were shown as TCID_50_/mL.

### RNA extraction and qRT-PCR analysis

Total viral RNA from cell culture supernatants was extracted by using the EasyPure® Viral DNA/RNA Kit (TransGen, Cat No. ER201-01 Beijing, China) according to the manufacturer’s instructions. Reverse transcription was performed to produce complementary DNA (cDNA) using Hifair 1st Strand cDNA Synthesis Kit (Yeasen Biotech, Cat No. 11123ES60, Shanghai, China), and then 2 μL cDNA was used as template for subsequent qPCR. The target N gene of GX_P2V isolate was amplified by qPCR using FastFire qPCR PreMix (Probe) (TIANGEN, Cat No. FP208-02, Beijing, China) according to the manufacturer’s instructions. Sequences of qRT-PCR primers are as follow: P2VF, TCTTCCTGCTGCAGATTTGGAT; P2VR, ATTCTGCACAAGAGTAGACTATGTA TCGT; and Probe, FAM-TGCAGACCACACAAGGCAGATGGGC-TAMRA. In addition, a standard plasmid was constructed through inserting a fragment amplified using P2VF and P2VR into a cloning vector pEASY-T1 (TransGen, Cat No. CT101-01 Beijing, China). Plasmids in a range of 10^2^ to 10^8^ DNA copies were used as template to generate a standard curve.

### Immunofluorescence assay (IFA) for viral growth

Confluent monolayers of BGM cells grown on coverslips in 12-well plates were incubated with GX_P2V at an MOI of 0.01 for two hours at 37°C in 5% CO_2_, then washed twice with PBS and cultured in RMPI 1640 with 2% FBS and HEPES (20 mM) at 37°C in 5% CO_2_. At indicated time points, the cells were washed twice with PBS and fixed with 4% paraformaldehyde for 20 min at room temperature, permeabilized by using 0.2% (vol/vol) Triton X-100 in PBS for 10 min, then blocked overnight with 3% bovine serum albumin (BSA) at 4°C. Then, cells were incubated with anti-sera (1:200, prepared in our laboratory) from GX_P2V-infected hamsters for two hours at room temperature. After being washed five times with PBS, cells were stained with a fluorescein isothiocyanate (FITC)-labeled goat anti-mouse IgG antibody (1:300, ZSGB-BIO, Cat No. ZF-0312, Beijing, China) for one hour at room temperature. Cells were washed five times and stained with 4',6-Diamidino-2-phenylindole (DAPI) (Sigma, Cat No. 10236276001, Germany) for 20 min. Finally, after being washed five times, cells on the coverslips were scanned and photographed using a fluorescence microscopy (Nikon, Japan).

### Golden hamster model of both intranasal and intragastrical infection

Specific pathogen-free, 6-week-old male golden hamsters were purchased from Charles River and maintained under specific-pathogen-free conditions. For intranasal infection, hamsters were infected with 10^4^ or 10^5^ TCID_50_ of GX_P2V in 50 μL RPMI 1640 containing 2% FBS after deep anesthesia with pentobarbital sodium. Equal volume of RPMI 1640 containing 2% FBS was set as mock-control group. The mock-infected group (n = 3) was monitored and sacrificed at 7 d.p.i. In the GX_P2V-infected group (n = 12 per group), three hamsters were sacrificed at each indicated time points (2, 5, 7 and 14 d.p.i). For intragastrical infection, hamsters were infected with 10^5^ TCID_50_ of GX_P2V in 100 μL RPMI 1640 containing 2% FBS. Equal volume of PBS was used to mock-infect the control group. Mock-infected hamsters (n = 4) were sacrificed at 14 d.p.i., while four hamsters of GX_P2V-infected (n = 20) were sacrificed at each indicated time points (2, 5, 7, 9 and 14 d.p.i).

After viral infection, various indexes were collected during 14-day experimental period. The body weight changes and clinical symptoms^35^ were recorded before the hamsters were humanly euthanized. Left lungs and bottom tracheas were placed in 4% paraformaldehyde and were stored at room temperature at least two days for observation of the pathological changes. The rest of the lungs and other organ tissues (stomach, duodenum, colon, tongue, trachea, esophagus, small intestine, feces) and sera samples were collected and frozen directly at −40°C for viral detection.

### Intranasal infection in both young and aged BALB/c mice

Specific pathogen-free, 6-week-old male young BALB/c mice and 48-week-old male aged BALB/c mice were obtained from SiPeiFu (Beijing). Young and old mice were randomly divided into two groups, respectively. Mice were anesthetized intraperitoneally (i.p.) with pentobarbital sodium and then infected intranasally with 10^5^ TCID_50_ of GX_P2V in 30 μL RPMI 1640 containing 2% FBS or PBS as control group. Mock-infected young (n = 5) or aged (n = 3) mice were sacrificed at 14 d.p.i., while five young or three aged GX_P2V-infected mice (n = 25 or 15, respectively) were sacrificed at each of the indicated time points (1, 3, 6, 9 and 14 d.p.i). Body weight and clinical symptoms were assessed periodically. Blood samples and lungs were collected regularly. Left lungs were fixed with 4% paraformaldehyde for determining histopathological changes. Right lungs were frozen directly at −40°C for analyzing the viral loads and titers.

### Measurement of viral RNA copies and virus titers in lungs and tracheas

Lung, trachea, stomach, duodenum, colon, tongue, esophagus, small intestine, and feces specimens were homogenized in 10 μL PBS per milligram of tissue for 60 s at 60 Hz 4 times using a Tissuelyse-24 system (Jingxin, Shanghai). Tissue homogenates were centrifuged at 5000 × *g* at 4°C for 5 min. Total viral RNAs from tissue supernatants were exacted by using EasyPure® Viral DNA/RNA Kit (TransGen, Cat No. ER201-01, Beijing, China) for qRT-PCR analysis as noted above. Infectious titers in tissue supernatants were detected and calculated by TCID_50_ assay as mentioned above. Virus titers were indicated as TCID_50_/mL.

### Histopathology of lungs and tracheas

The lungs and tracheas were grossly examined and photographed before freezing and fixing. The left lungs and bottom tracheas were fixed in 4% paraformaldehyde and embedded in paraffin. The paraffin sections were prepared for H&E staining. The histopathology of the lungs and tracheas tissues was detected by microscopy (Nikon, Japan). The histopathological results were analyzed by professional pathologists.

### Anti-sera preparation from hamsters

Hamsters were intranasally challenged with two doses (1 d.p.i and 21 d.p.i) of 10^5^ TCID_50_ GX_P2V. Sera were collected at 42 d.p.i for analysis of antibody responses by immunofluorescence assay (IFA), ELISA, and pseudovirus neutralization titer assay (pVNT). Anti-sera were heated at 56°C for 30 min before use.

### Immunofluorescence assay for cross-immunity responses

The full-length spike protein gene of SARS-CoV-2 (accession number NC_045512.2) and GX_P2V (accession number MW532698) were codon optimized, synthesized, and cloned into pcDNA3.1 vector by Beijing BioMed Gene Technology. Confluent monolayers of 293T cells grown on coverslips in 12-well plates overnight were transfected with plasmids containing codon-optimized spike genes of SARS-CoV-2 and GX_P2V by using liposome (TransGen, Cat No. FT201-02, Beijing, China) according to the instructions. After five hours, the cells were washed twice with PBS and cultured in DMEM with 2% FBS at 37°C in 5% CO_2_. After 36 h post-transfection, cells were fixed, permeabilized, and blocked. Then, cells were incubated with sera from GX_P2V-infected hamsters (1:200), convalescent COVID-19 patients (1:200), mock-infected hamsters, or healthy donors. After being washed five times with PBS, cells were stained with secondary antibodies: fluorescein isothiocyanate (FITC)-labeled goat anti-mouse IgG (1:300, ZSGB-BIO, Cat No. ZF-0312, Beijing, China) or goat anti-human IgG (1:300, ZSGB-BIO, Cat No. ZF-0308, Beijing, China), accordingly. Following five washes with PBS, cells were stained with 4',6-Diamidino-2-phenylindole (DAPI) (Sigma, Cat No. 10236276001, Germany) for 10 min. Finally, after being washed five times, cells on the coverslips were analyzed using fluorescence microscopy (Nikon, Japan).

### ELISA binding assay

SARS-CoV-2 RBD, S1 and S1+S2 (ECD) antibody titer assay kits (Sino Biological, Cat No. KIT002, KIT003, KIT004, respectively, Beijing, China) were purchased. Experiments were carried out following the manufacturer’s instructions with minor modifications. All sera were diluted with dilution buffer at a ratio of 1:100. Patient’s sera were tested according to the instructions. For testing sera from hamsters and BALB/c mice, the secondary antibodies were replaced by HRP-goat anti-mouse IgG (1:5000, ZSGB-BIO). Plates were read at 450 nm wavelength on a synergy H4 hybrid reader (Biotek, USA).

### Pseudotyped virus neutralization assay

The VSV-based SARS-CoV-2 pseudovirus was kindly provided by Prof. Youchun Wang (National Institutes for Food and Drug Control). The SARS-CoV-2 pseudovirus was used as seed stock and combined with the GX_P2V spike protein expression plasmid to generate VSV-based GX_P2V pseudovirus in 293T-hACE2 cells. The pseudovirus based neutralization assay procedure was conducted as previously described with minor modifications^36^. Briefly, sera samples were diluted in five three-fold serial dilutions and incubated with 1000 TCID_50_ of pseudovirus in 96-well plates, making the initial dilution 1:30. After one h of incubation at 37◻, cells (2.5×10^4^ cells/well) were added and incubated for another 24 hours at 37◻ in a CO_2_ incubator. Cells were lysed by adding luciferase substrate (PerkinElmer, USA). Luciferase activity was then measured using a synergy H4 hybrid reader (Biotek, USA). The 50% neutralization titer (NT_50_) was calculated for each individual serum sample by using the Reed-Muench method.

### Statistical analysis

The statistical analyses were performed with GraphPad Prism version 9.0 software. The differences of two groups were analyzed by two-tailed Student *t*-test. The *p*-values of less than 0.05 (*p* < 0.05) were considered statistically significant.

## Acknowledgements

We thank Harlan D. Caldwell for critical review of the manuscript. This research was supported by NSFC-MFST projects (China-Mongolia) (No. 3211101856 and 31961143024), Inner Mongolia Key Research and Development Program (No. 2019ZD006), Funds for First-class Discipline Construction (No. XK1805 and XK1803-06), National Key Research and Development Program of China (No. 2018YFA0903000, 2020YFC2005405, 2020YFA0712100, 2020YFC0840805, 19SWAQ06, 20SWAQX27 and 20SWAQK22), Fundamental Research Funds for Central Universities (No. BUCTRC201917 and BUCTZY2022).

## Author contributions

We contributed to the work as follows. L.S., Y.T. and P.M. designed and supervised research. S. Lu, S. Luo, C.L., M.L., J.L., X.A., Z.L., Y.Y., J.H., and H.F. conducted experiments. P.M. and L.S. performed analyses. S. Lu, S. Luo and L.S. wrote the paper. All of us reviewed and approved the manuscript.

